# Members of Venezuelan Equine Encephalitis Complex entry into host cells by clathrin-mediated endocytosis in a pH-dependent manner

**DOI:** 10.1101/2022.03.11.483939

**Authors:** Lucia Maria Ghietto, Pedro Ignacio Gil, Paloma Olmos Quinteros, Emiliano Gomez, Franco Martin Piris, Patricia Kunda, Marta Contigiani, Maria Gabriela Paglini

**Affiliations:** Instituto de Virología “Dr. JM Vanella”, Facultad de Ciencias Médicas, Universidad Nacional de Córdoba, Córdoba, Argentina; Instituto de Investigación Médica Mercedes y Martín Ferreyra, INIMEC-CONICET-Universidad Nacional de Córdoba, Córdoba, Argentina; Centro de Investigación en Medicina Traslacional “Severo Amuchástegui” (CIMETSA), Instituto Universitario Ciencias Biomédicas Córdoba (IUCBC), Naciones Unidas 420, Córdoba, Argentina

**Keywords:** Pixuna virus, Rio Negro virus, Alphavirus, entry, endosomes

## Abstract

Pixuna virus (PIXV) and Río Negro virus (RNV) are mosquito-borne alphaviruses belonging to the Venezuelan Equine Encephalitis (VEE) complex, which includes pathogenic epizootic and enzootic subtypes responsible for life-threatening diseases in equines. Considering that the first steps in viral infection are crucial for the efficient production of new progeny, the aim of this study was to elucidate the early events of the replication cycle of these two viruses. To this end, we used chemical inhibitors and the expression of dominant-negative constructs to study the dependence of clathrin and endosomal pH on PIXV and RNV internalization mechanisms. We demonstrated that both viruses are internalized primarily via clathrin-mediated endocytosis, where the low pH in endosomes is crucial for viral replication. Contributing knowledge regarding the entry route of VEE complex members is important to understand the pathogenesis of these viruses and also to develop new antiviral strategies.

## 1. Introduction

The Venezuelan equine encephalitis (VEE) antigenic complex belongs to the Togaviridae family, which include the Alphavirus genus. They constitute a group of arboviruses responsible for human and veterinary disease outbreaks around the world ^1^, they share similar genetic characteristics and can be defined by their broad antigenic cross-reactivity ^2^. The VEE complex comprises New World viruses that cause encephalitis in humans and horses and are classified in six antigenic subtypes divided into epizootic (IAB and IC) and enzootic subtypes ^3^. One of the members of the VEE complex is Pixuna virus (PIXV), New World alphavirus subtype IV ^4^. PIXV was first isolated from Anopheles nimbus mosquitoes in Brazil in 1961 ^5^. It has been detected in Argentina ^6,7^ and it is a non pathogenic member of the complex. Another member is Rio Negro virus (RNV), subtype VI, that circulates in Argentina ^6,7^ and which has been isolated from rodents ^8^. In 1989, RNV caused an outbreak of acute febrile illness in humans, which was originally attributed to dengue virus due to the similarity of the symptoms they both cause ^8^. Enzootic subtypes, including PIXV and RNV, are also important for human health because they can cause febrile illnesses with clinical symptoms similar to those caused by dengue or influenza (headache, chills, fever, myalgia, retroocular pain) ^9^.

Viruses use the host cell machinery for entry, replication, transport and release of the viral progeny. The first steps in viral infection are crucial for the efficient production of new progeny. The different stages of VEE complex enzootic variants replication cycle are still poorly understood. Notwithstanding, studies on alphaviruses have demonstrated that after binding to receptors on the plasma membrane they enter into the cell via clathrin-mediated endocytosis and fusion with endosomal membranes ^10–12^. However, the mechanisms concerning the cell biology of alphaviruses infection are still unclear. Some authors have described other pathways for cell penetration, including caveolae-dependent endocytosis ^13^, a direct delivery through a putative pore at the plasma membrane ^14^ or macropinocytosis-mediated uptake ^15^. In this sense, and considering that the entry route of different enzootic variants of the VEE complex remain poorly explored, it is important to describe the mechanisms of entry and uncoating of these viruses. Therefore, the aim of the present research was to elucidate the early events of the replication cycle of PIXV and RNV, two enzootic members of the VEE complex. To this end, we used chemical inhibitors and the expression of dominant-negative constructs to study the dependence of clathrin and endosomal pH on PIXV and RNV internalization mechanisms. Agents that disrupt clathrin-dependent receptor-mediated endocytosis such as Sucrose (SCR) and Chlorpromazine (CPZ) were used. SCR was necessary to generate a hypertonic medium to disturb the formation of clathrin vesicles on the cell membrane ^16–21^. CPZ was used as it prevents clathrin-coated pit formation on the plasma membrane by promoting clathrin lattice assembly on endosomal membrane ^22–25^. The lysosomotropic agents, Glucosamine (GlcN) and ammonium chloride (NH_4_Cl) were used to elevate acidic organelles pH owing to the presence of their amino group ^10,26–30^. The above mentioned compounds have been widely used to decrease viral replication in several enveloped viruses ^31–35^. We demonstrate that both viruses are internalized primarily via clathrin-mediated endocytosis, where the low pH in endosomes is crucial for viral replication. The results reported here show new insights regarding VEE complex members entry route and infection. In addition, these findings provide a better understanding of the molecular mechanisms underlying their replication cycle pointing to attractive targets for the development of new strategies to prevent infections.

## 2. Results

### 2.1. Clathrin is required for PIXV and RNV entry into Vero cells

Considering that the ability of alphaviruses to enter into the host cell via alternative pathways has already been reported ^13–15,36^, we evaluated the dependence of PIXV and RNV entry into Vero cells by clathrin-mediated endocytosis. To this end, we first employed chemical inhibitors, widely used for clathrin-mediated endocytosis, such as SCR ^16–21^ and CPZ ^22–25^. According to the literature and to the possible drug-induced cytotoxic effects, cell culture viability in the presence of MTT was assayed at different concentrations of these inhibitors (Figure 1A). Thus, we defined 0.2 M for SCR and 25 μM for CPZ as the experimental conditions (cell viability ≥ 90%). Besides, the inhibitory action of both drugs on clathrin-mediated endocytosis was effectively assessed determining TRITC-transferrin internalization reduction. The fluorescence inside the cell cytoplasm observed in cultures treated with SCR or CPZ was significantly lower (64.6% and 55.9%, respectively; p < 0.001) compared to control cells (100%, Figure 1B). After PIXV or RNV internalization, in the presence of drugs, we collected the supernatants at 4, 8 and 24 hours postinfection (hpi) and quantified extracellular infectious particles by plaque assay (plaque-forming unit, PFU/ml). Infected cultures with PIXV (multiplicity of infection: MOI 0.1) and treated with 0.2 M SCR or 25 uM CPZ showed a significant decrease in PUF/ml of about 2 log_10_ and 1-2 log_10_ at 8 and 24 hpi, respectively. In the same way, infected cultures with RNV (MOI 0.1) and treated with both drugs displayed a significant reduction of virus particles in the extracellular medium of about 1-2 log_10_ and 2 log_10_ at 8 and 24 hpi, respectively (Figure 1C). As we expected, no differences in extracellular virus yield was observed at the end of the eclipse phase (4 hpi) ^37^ for either viruses. Additionally, cell cultures were treated and infected with PIXV or RNV (MOI 10). At 8 hpi, cultures were fixed and processed for immunofluorescence in order to calculate the percentage of infected cells. In the representative image, the culture shows a visible reduction of green fluorescence labeling after treatment with both inhibitors in comparison to mock-treated cells (Figure 1D). Cultures treated with SCR or CPZ showed a significant reduction in the percentage of PIXV (67.83 ± 3.68 and 56.15± 25.85, respectively; p < 0.001) and RNV (51.17 ±18.49 and 21.49 ± 16.7 respectively; p < 0.001) infected cells with respect to control (100%). Additionally, we observed significant differences between both drugs in cultures infected with RNV (p < 0.001) (Figure 1E).

**Figure 1.**
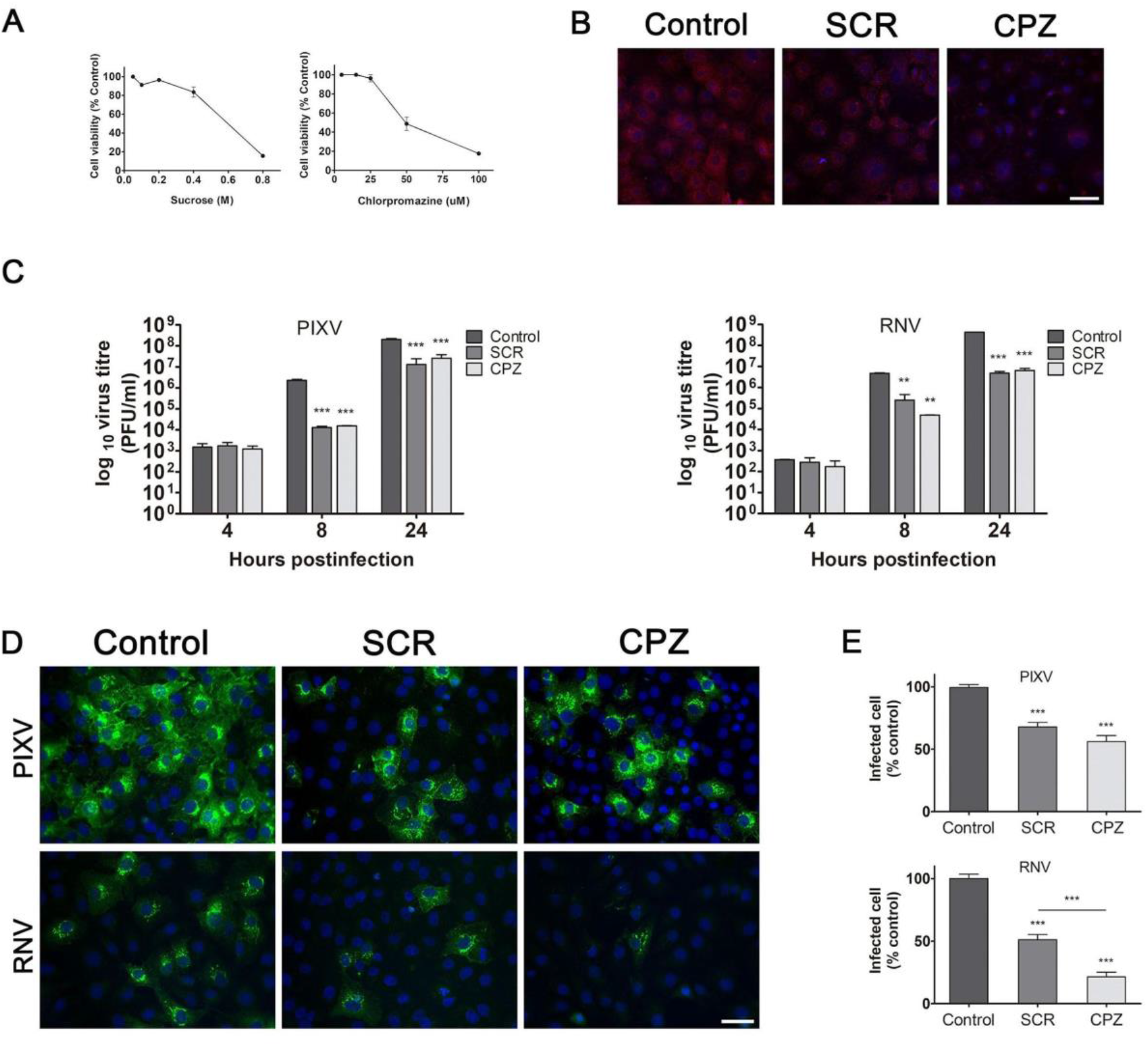
Assays with inhibitors targeting clathrin-mediated endocytosis of PIXV and RNV entry. **A.** MTT assay to determine SCR and CPZ cytotoxic concentrations. **B.** Representative images of Vero cells treated with 0.2 M SCR or 25 μM CPZ and incubated with TRITC-transferrin (red); nuclei (blue). **C.** Cells were treated with 0.2 M SCR or 25 μM CPZ and infected with PIXV or RNV at MOI 0.1. Supernatants were collected at 4, 8 and 24 hpi and extracellular infectious particles were quantified by plaque assay (plaque-forming unit, PFU/ml). **D**. Representative images of cultures treated with 0.2 M SCR or 25 μM CPZ and infected with PIXV or RNV (green) at MOI 10 and fixed at 8 hpi; nuclei (blue). **E**. Percentage of infected cells with PIXV or RNV against control (100% value). Data represent the mean ± SD from 3 independent experiments. Significance was calculated by one-way ANOVA test, followed by Tukey post-hoc analysis to enable specific group comparisons, with p value ≤ 0.05 considered as statistically significant. **p < 0.01; ***p < 0.001. Scale bar: 20 μm.

To support the participation of the clathrin pathway in PIXV and RNV entry, we employed wild type (WT) and dominant negative (DN) mutant constructs of the clathrin-coat associated cellular protein Eps15, a required adaptor for clathrin-mediated endocytosis ^38^. Vero cells were transfected with plasmids expressing Eps15-WT or Eps15-DN; and their functionality was confirmed by incubation of transfected cells with TRITC-transferrin. Eps15-DN expression significantly reduced the fluorescence intensity of the cell cytoplasm (46.29 %; p < 0.001, n=15-17 cells for each condition) compared to cells expressing Eps15-WT (100 % value) (Figure 2A). Transiently transfected cells with either Eps15-WT or Eps15-DN were then infected with PIXV or RNV (MOI 10). The effect of mutant proteins expression on the internalization of both viruses was analyzed by immunofluorescence at 8 hpi (Figure 2B). No significant differences were found in the percentage of infection in all cultures transfected under either condition (24.51 ± 10.53 Eps15-WT; 19.18 ± 7.75 Eps15-DN for PIXV and 14.74 % ± 2.38 Eps15-WT; 12.34 ± 5.35 Eps15-DN for RNV). Quantification of the transfected/infected cells showed a significant reduction in infection of about 60% (p < 0.001) for the Eps15-DN condition relative to the Eps15-WT for both viruses (Figure 2C). Taken together, these results provide strong evidence that clathrin plays an important role for both PIXV and RNV entry into host cells.

**Figure 2.**
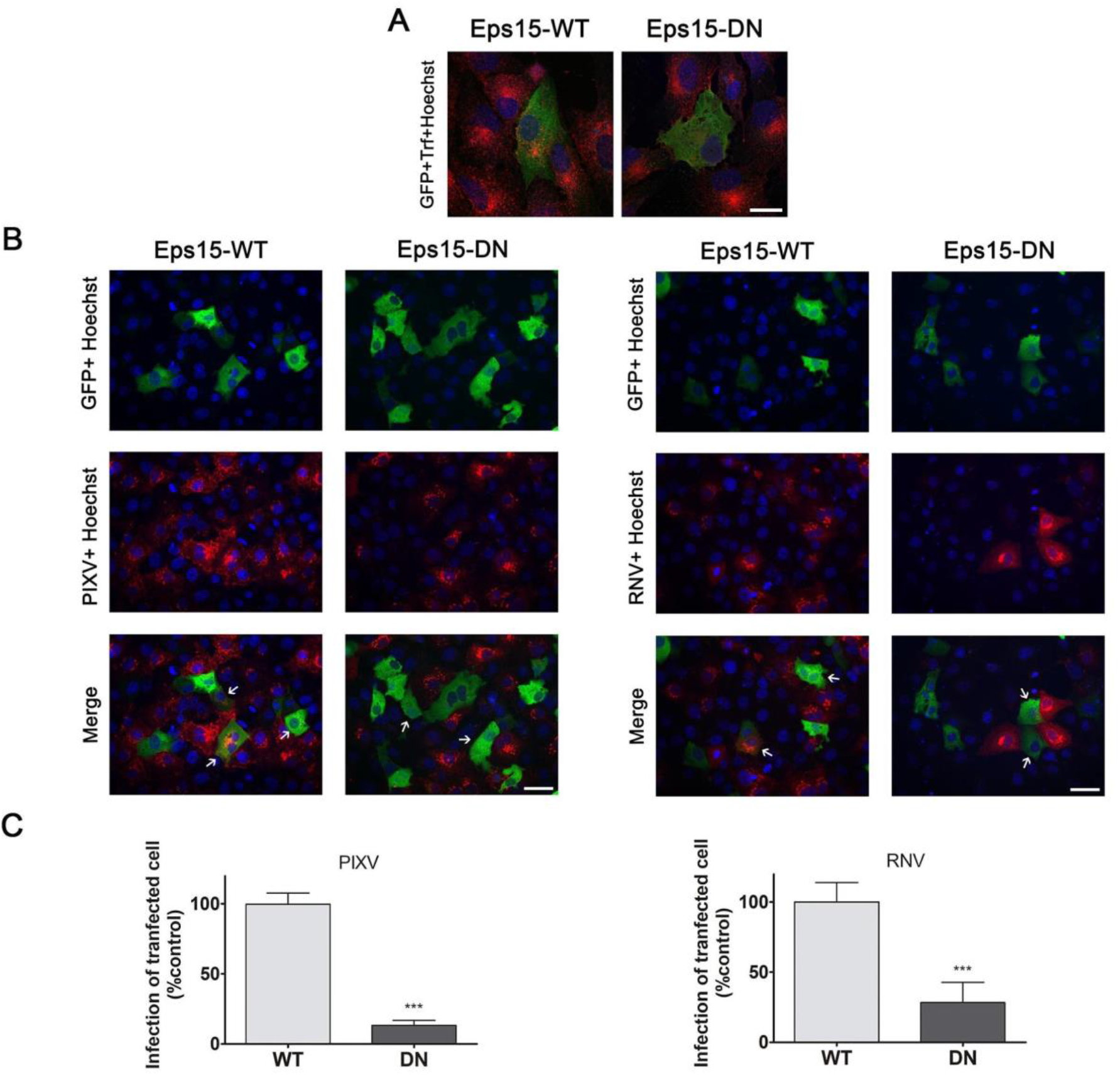
Inhibition of PIXV and RNV entry by transient expression of the Eps15 dominant negative mutant construct. **A**. Representative images of cells transfected with plasmids expressing Eps15-WT or Eps15-DN (green) and incubated with TRITC-transferrin (red); nuclei (blue). **B.** Immunofluorescence of transiently transfected cells with plasmids expressing Eps15-WT or Eps15-DN (green) and infected with PIXV or RNV (red) at MOI 10 after 8 hpi (arrow); nuclei (blue). **C.** Percentages of transfected and infected cells. More than 300 transfected cells were screened in each case. Data represent the mean±SD from 3 independent experiments. Significance was calculated by Student’s t test, with p value ≤ 0.05 considered as statistically significant. ***p < 0.001. Scale bar: 20 μm.

### 2.2. PIXV and RNV infection requires active endosomal acidification

Endosomal vesicles may play a crucial role in virus infection, providing the appropriate condition for conformational changes in envelope viral proteins, resulting in the formation of a fusion pore and release of the viral nucleocapsid into the cytosol ^39^. Considering that the endocytic pathway is highly regulated and involves several intermediate organelles, it is important to examine the fate of viral proteins after their internalization in early and late endosomes. The Rab family of small GTPases is essential for membrane trafficking and the classification of endocytic loads into specific subcellular compartments ^40^. In this sense, we performed a quantitative colocalization analysis of PIXV and RNV with Rab5-GFP and Rab7-GFP, as early and late endosome markers, respectively ^41^ (Figure 3A). Using Pearson’s colocalization coefficient ^42^, we observed significant colocalization PIXV and RNV proteins with Rab5-GFP and Rab7-GFP (Figure 3A). After 8 hpi, a great coefficient of colocalization was found for both endosomal compartments and the viral proteins (PIXV: r= 0.68 ± 0.11 for Rab5 and r= 0.78 ± 0.06 for Rab7; RNV: r=0.69 ± 0.1 for Rab5 and r= 0.75± 0.07 for Rab7) (Figure 3B). These results show that PIXV and RNV associate to early and late endosomes at the early stage of the infectious cycle.

**Figure 3.**
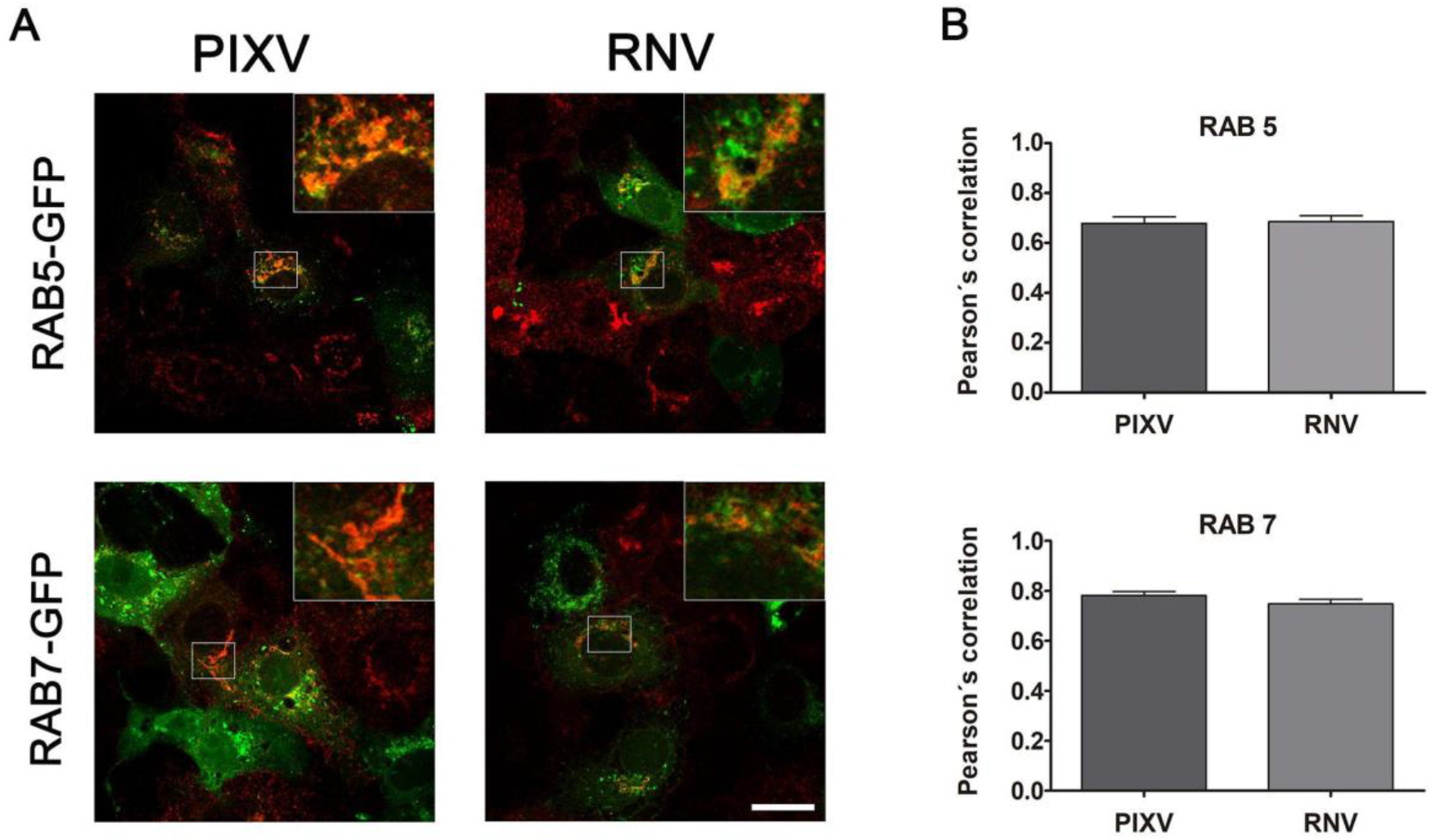
Colocalization of PIXV and RNV with Rab5 and Rab7. **A.** Immunofluorescence of transiently transfected cells with plasmids expressing Rab5-GFP or Rab7-GFP (green) and infected with PIXV or RNV (red) at MOI 10. **B.** Colocalization analyses for Rab5-GFP or Rab7-GFP expression with PIXV or RNV were expressed as Pearson’s correlation coefficient. The plots show the mean of Pearson’s correlation coefficients ± SD, n=10-12 cells for each condition from three independent experiments. Scale bar: 20 μm.

To study the low pH-dependence of PIXV and RNV replication, we first evaluated the effect of two lysosomotropic agents, GlcN and NH_4_Cl, inhibitors of endosomal vesicles acidification ^10,26–33^, in Vero cells. Cell culture viability by MTT was assayed at different concentrations of either of these two inhibitors (Figure 4A). Thus, we defined 15 mM GlcN and 25 mM NH_4_Cl as the experimental conditions (cell viability ≥ 90%). To assure the effect of both compounds in intracellular acidic vesicles, we pretreated Vero cells with the drugs plus acridine orange. Cultures treated with NH_4_Cl showed significant differences in the fluorescence intensity of the cells cytoplasm (56.3 %; p < 0.001) compared to controls (100%). However, this effect was not observed in culture cells treated with GlcN, where acridine orange fluorescence inside the cell cytoplasm did not display significant differences (87.1%) compared to the control condition (Figure 4B). We then performed the cytoplasmic accumulation assay with neutral red, a very sensitive probe for lysosomal microenvironments and a biological pH indicator ^43^. In figure 4C, a 40% and 80% inhibition of the neutral red signal can be observed in cultures treated with GlcN and NH_4_Cl, respectively, compared to untreated cells. Therefore, the participation of endosomal pH in PIXV and RNV replication in Vero cells was analyzed by extracellular virus titration. Viral progeny production of PIXV and RNV was reduced by about 3 log_10_ and 4 log_10_ in cultures incubated with GlcN or NH_4_Cl respectively, at 8 and 24 hpi, compared to control cultures (Figure 4D). As we expected, no differences in extracellular virus yield was observed at the end of the eclipse phase (4 hpi) ^37^ for either virus. The fluorescence pattern of intracellular PIXV proteins observed in control cultures was substantially reduced in cells treated with GlcN or NH_4_Cl. In cultures infected with RNV, fluorescence was highly reduced when cells were treated with both compounds (Figure 4E). A reduction of 50% was detected in the number of cells expressing PIXV proteins in cultures treated with GlcN and approximately 100% in those treated with NH_4_Cl compared to untreated infected ones. Interestingly, the percentage of RNV infected cells was practically zero in cultures treated with both lysosomotropic agents (Figure 4F).

**Figure 4.**
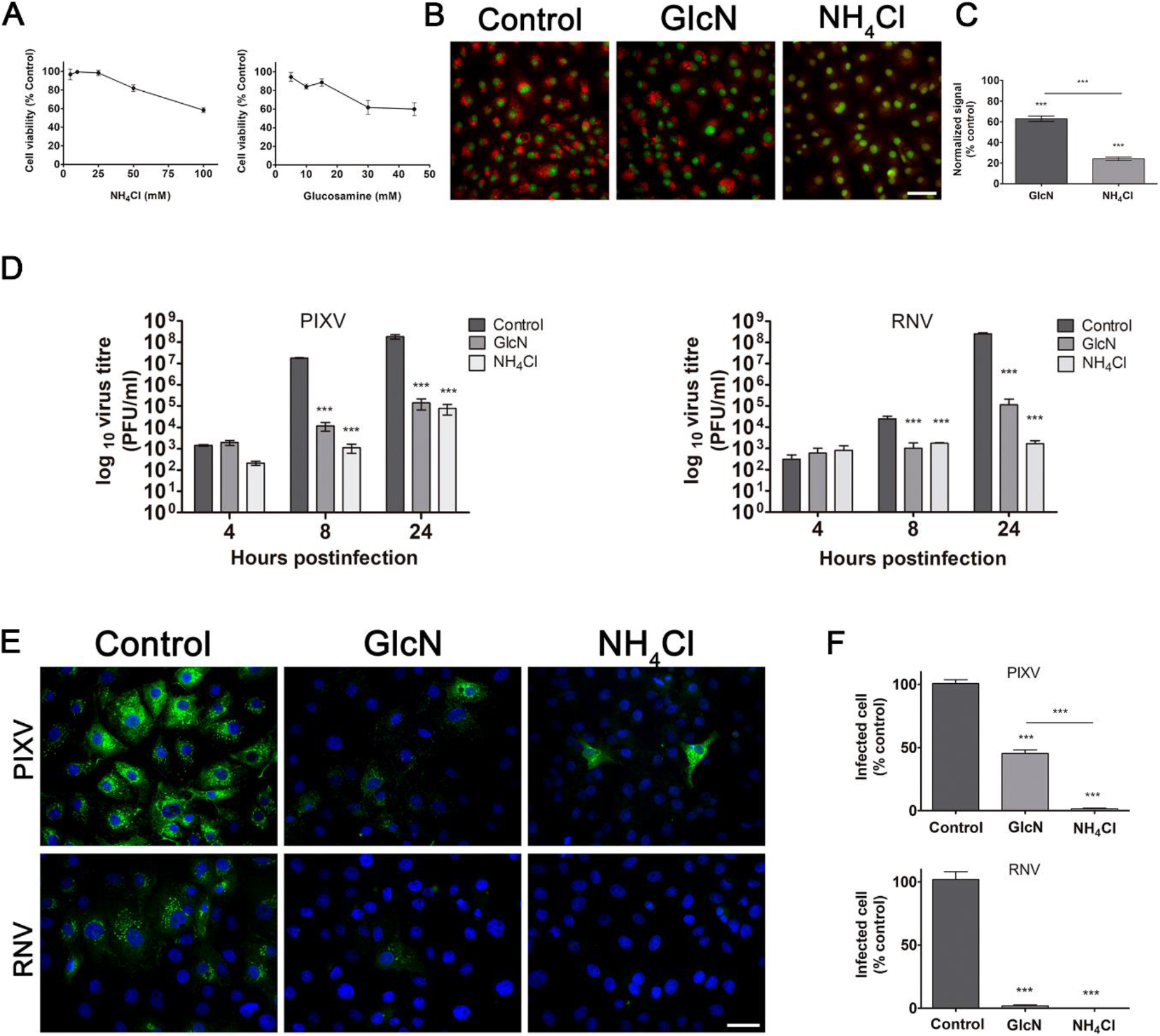
Effect of lysosomotropic agents on PIXV and RNV internalization. **A.** MTT assay to determine GlcN and NH_4_Cl cytotoxic concentrations. **B.** Representative images of Vero cells treated with 15 mM GlcN or 25 mM NH_4_Cl and incubated with acridine orange (red); nuclei (green) **C.** Vero cells incubated with 15 mM GlcN or 25 mM NH_4_Cl and neutral red (NR). Percentages of inhibition were calculated by absorbance compared to control cultures. **D.** Cells were pretreated with 15 mM GlcN or 25 mM NH_4_Cl and infected with PIXV or RNV at MOI 0.1 (green). Supernatants were collected at 4, 8 and 24 hpi and extracellular infectious particles were quantified by plaque assay (plaque-forming unit, PFU/ml). **E**. Representative images of cultures treated with 15 mM GlcN or 25 mM NH_4_Cl and infected with PIXV or RNV (green) at MOI 10 and fixed at 8 hpi; nuclei (blue). **F**. Percentage of infected cells with PIXV or RNV against controls (100% value). Data represent the mean ± SD from 3 independent experiments. Significance was calculated by one-way ANOVA test, followed by Tukey post-hoc analysis to enable specific group comparisons, with p value ≤ 0.05 considered as statistically significant. *p < 0.05; **p < 0.01; ***p < 0.001. Scale bar: 20 μm.

## 3. Discussion

Alphaviruses constitute a conserved group of arboviruses responsible for human and veterinary disease outbreaks around the world ^1^. The VEE complex comprises varieties of enzootic and epizootic viruses. The epizootic strains emerge periodically in regions of Central America and northern areas of South America, causing outbreaks that involve humans and horses, with high morbidity and mortality rates ^1,2^. In particular, Venezuelan Equine Encephalitis virus (VEEV) is the most important zoonotic pathogen, with several reported outbreaks in South and Central America ^44^. Alphaviruses are enveloped viruses that, in general, penetrate into the cells by different ways: by fusion with the cell plasma membrane or by receptor-mediated endocytosis; a pathway that usually relies on endocytic vesicles low pH. Different endocytic pathways have been characterized over the past decade, among which the clathrin-mediated endocytosis is one of the entry routes most commonly used by many enveloped viruses ^45^. Moreover, this is the most widely accepted mechanism for alphaviruses entry to the host cell and for the subsequent fusion between the viral envelope and the endosomal membrane ^46^. However, some reports showed alternative pathways, such as early endosomes and caveolae-derived vesicles for Mayaro virus in Vero cells ^13^, the direct delivery through a putative pore at the plasma membrane for Sindbis virus ^14^, or macropinocytosis for Chikungunya virus (CHIKV) entry into human muscle cells ^15^. To investigate whether in Vero cells two members of the VEE antigenic complex use clathrin in the endocytic pathway, various systematic approaches were used. These include pharmacological inhibition and overexpression of Eps15 WT and DN mutant plasmids. Our results suggest that infection of target cells depends, in part, on clathrin-mediated endocytosis as endocytic inhibitors only resulted in partial block of PIXV and RNV infection. This supports the hypothesis that several pathways are used by alphaviruses to facilitate their entry into target cells ^47^. It is well known that actin assembly plays an essential role in several processes leading to endocytosis, including the internalization of clathrin-coated vesicles ^48^. In agreement with previous literature, our work supports the notion that PIXV and RNV use clathrin-mediated endocytosis as the major/principal pathway to enter into the host cell. In this sense, previous results from our laboratory demonstrated that PIXV relies on a dynamic actin network during the early steps of viral replication. This indicates that specific interactions with cytoskeletal proteins are necessary for the normal development of viral replication ^37^. Different members of the alphavirus genus have been reported to enter host cells using this pathway ^47^. Nevertheless, mutants of Eps15 and Rab5, as well as endocytic inhibitors only resulted in partial block of CHIKV infection, supporting the hypothesis that several pathways are hijacked by the virus to facilitate its entry into target cells ^32^. In line with this idea, Lee and collaborators demonstrate that CHIKV entry into human muscle cells is mediated by macropinocytosis ^15^. Besides, new findings show that VEEV and Western equine encephalitis virus exploit the caveolin-1-mediated transcytosis via to enter the central nervous system ^49^, providing alternative pathways for virus entry into different types of cells. For example, as described for Influenza virus, clathrin-endocytosis and macropinocytosis can operate simultaneously in HeLa, A549 and other non-human cells ^50^. Considering that PIXV and RNV entry depends only in part on clathrin-mediated endocytosis, it would be interesting to study alternative pathways that might be used to facilitate their entry into target cells.

In alphaviruses, as well as in other viruses, the progressive pH decrease in the maturing endocytic pathway triggers events in the viral envelope proteins resulting in viral uncoating. The E2 envelope protein mediates viral entry by attachment to cellular receptors. Then, the low-pH environment triggers a sequence of events starting with the dissociation of the E1-E2 dimer and followed by conformational changes in the E1 membrane fusion protein. This sequence of events results in the formation of a fusion pore and the release of the viral ARN into the cytosol ^47,51^. Almost all viruses that are internalized by the cell via endocytosis are first delivered to Rab5-positive early endosomes before being targeted to Rab5/Rab7-positive maturing endosomes, Rab7-positive late endosomes, and lysosomes ^52^. Our results show that PIXV and RNV proteins colocalize with both, early and late endosomes, suggesting that these viruses enter the endocytic pathway of the host cell and require trafficking as far as early and late endosomes to initiate an infectious cycle. Previous studies on alphaviruses revealed that Semliki Forest virus fused with early endosomes, and VEEV was found to fuse with maturing and/or late endosomes ^10–12^. In CHIKV, membrane fusion was observed predominantly with Rab5-positive endosomes and often occurred within 40 s after delivery to endosomes ^53^. To further confirm our observations, we employed lysosomotropic agents to interfere with endosome acidification that are extensively used as an assay for alphaviruses entry ^10,54^. As expected, PIXV and RNV required a progressive pH reduction in endosomes in order to release the viral nucleic acid into the cytosol.

In conclusion, this study demonstrates, for the first time, that PIXV and RNV partially enter via clathrin-dependent endocytosis into Vero cells. Then, both viruses require transport to early and late endosomes, where the acidic pH-dependent step results in the release of PIXV or RNV RNA into the cytoplasm for a successful infection (Supplementary Fig.1). This study contributes to our understanding of the entry process in enzootic subtypes from the VEE complex. Considering that arthropod borne viruses are significant sources of disease in man and domestic animals, this research extends the knowledge of virus entry into host cells and provide a better understanding of the molecular mechanisms underlying the viral internalization process. Ultimately, this knowledge would be useful to develop potent antivirals that could block the virus at the early stages of infection.

## 4. Materials and Methods

### 4.1. Cell culture and viruses

Vero cells (Vero 76, ATCC CRL 1587) were grown in Dulbecco’s modified Eagle’s medium (DMEM) (GIBCO, USA) with 10% fetal calf serum (FCS), Gentamicin (50 μg/ml), at 37 °C, 5% CO2 and 98% humidity. The PIXV strain BeAr 35645, isolated from *Anopheles nimbuspor* in Brazil ^5^, was propagated in Vero cells as previously described ^55^ and stocks were stored at 2 × 10^8^ plaque-forming units (PFU/ml). The RNV strain AG80-663, isolated from *Culex delpontei* in 1980 ^56^, was propagated in Vero cells, according to Pisano *et al*., 2012 ^57^ and stocks were stored at 7 × 10^8^ PFU/ml. Virus inoculum (PFU/ml) was determined by a standard plaque assay.

### 4.2. Viral infection

The protocol of synchronized infection has been previously described by Gil *et al*., 2017 ^37^. Briefly, Vero cells were plated and incubated at 37 °C, 5% CO2 and 98% humidity. Confluent monolayers of Vero cells were infected at different multiplicity of infection (MOI); MOI 0.1 for standard plaque assays and MOI 10 and for immunofluorescence assays. Virus adsorption was performed at 4 °C per 1 h. After this first incubation, the inoculum was removed and monolayers were washed with PBS at 4 °C to eliminate any free virus particle, then DMEM with 1% FCS was added and the infected cultures were incubated at 37 °C, 5% CO2 and 98% humidity for 8, 12 or 24 h, depending on the experiment. The cytopathic effect was evaluated under direct visualization by phase contrast microscopy at low magnification using an Olympus IX81 microscope.

### 4.3. Virus titration by plaque assay

Infected Vero cell monolayers with serial dilutions of each culture supernatant collected from the different assays were overlaid with 1.5% ultra-pure agarose (Gibco BRL) in 4% DMEM 2x. Two days later, the cells were fixed with 10% formaldehyde in water and the overlaid medium was removed. The monolayers were stained with 1% crystal violet in water and the plaques were counted. Infectivity titers were expressed as PFU/ml. All the experiments with viruses were performed in a laminar flow hood under biosafety level 2 conditions.

### 4.4. Drug treatments

Sucrose (SCR) was used to generate a hypertonic medium to disturb the formation of clathrin vesicles on the cell membrane. Under this condition, clathrin-dependent endocytosis is inhibited ^16–21^. Chlorpromazine (CPZ) was used in this study as an inhibitor of clathrin-mediated endocytosis. CPZ specifically inhibits clathrin-coated pit formation on the plasma membrane ^22–25^. The lysosomotropic agents, Glucosamine (GlcN) and ammonium chloride (NH_4_Cl) were used to elevate the pH of acidic organelles owing to the presence of its amino group ^10,26–30^. Those compounds have been widely used to decrease viral replication in several enveloped viruses ^31–33^. All compounds were purchased from Sigma-Aldrich and were dissolved in phosphate-buffered saline (PBS) to prepare master-stocks for storage.

To evaluate the efficiency of blockade of each endocytic pathway by the different inhibitors, uptake studies with TRITC-transferrin (Transferrin From Human Serum, Alexa Fluor™ 647 Conjugate #T23366, Thermo Fisher Scientific Inc., Waltham, Massachusetts, USA) for SCR and CPZ, neutral red and acridine orange for NH_4_Cl and GlcN were performed. We used those strategies because transferrin and its receptor are taken into cells through clathrin-mediated endocytosis, a mechanism that has been extensively characterized ^58^. Neutral red and acridine orange are dyes commonly used as probes to measure the internal pH in acidic vesicles ^59^. Untreated cells were used as control.

Cells either treated with inhibitors or not, were incubated with 15 μg/ml TRITC-transferrin or 100 μg/ml acridine orange (A6014, Sigma) in serum free medium during 15 min at 37°C. Then, cells were processed for visualization in a fluorescence microscope as described below. For the quantification of fluorescence intensity of TRICT-transferrin or orange acridine, images of 5 microscope fields, selected at random, were acquired per condition in each independent experiment. Using the FIJI software (https://fiji.sc/), fluorescence intensity was calculated and data were normalized against fluorescence of controls (100% value). Background intensity was subtracted from images prior to quantification ^60^.

To determine inhibition by lysosomotropic agents, we incubated cultured cells with drugs at a selected concentration plus 50 mg/ml neutral red (NR), as a biological lysosomal pH indicator. Cells were seeded in 100 μl per well of transparent 96-well microtiter plates. Following drug treatments, the cell culture medium was replaced with medium containing 50 μg/ml NR dye and drugs. Following incubation for 3 h at 37 °C and 5% CO2 in a humidified environment and two washes with PBS, intracellular NR dye was extracted using a destaining solution (50% ethanol, 1% acetic acid and water) and absorbance was measured at 540 nm using a spectrophotometer (Biotek, Elx800). Percentage of inhibition of each drug was calculated normalizing data against the negative control absorbance (100% value).

To evaluate the effect of different drugs on PIXV and RNV infection, Vero cell monolayers were seeded into 6-well plates or 24-well plates and pretreated with drugs, as listed before, for 30 min at 37 °C. Synchronized infection of PIXV or RNV was achieved by infecting the cells at MOI of 0.1 or 10 at 4 °C for 1 h in the presence of the compounds. Then, the inoculum was removed and DMEM was added with different inhibitors throughout all designed experiments. In cultures infected at MOI 0.1, the supernatant was collected at 4, 8 and 24 hpi to determine extracellular viruses by a titration assay. Monolayers infected with MOI 10 were fixed at 8 hpi to proceed to an immunofluorescence assay to determine intracellular viruses.

### 4.5. Viability assay

Cell viability under the different drug treatments were assessed by employing 3-(4 5-dimethylthiazol-2-yl)-5-(3-carboxymethoxyphenyl)-2-(4-sulfophenyl)-2h-tetrazolium (MTT). Briefly, Vero cells were seeded at the density of 1 × 10^6^ cells/well in a 96-well plate. After 24 h, cells were treated with drugs at different concentrations for 24 h, untreated cells served as control. After treatment, cells were washed with PBS and 100 μl MTT assay was mixed with 900 μl PBS into each well, followed by incubation at 37 °C for 2 h. The medium was then removed, washed, and 1 ml DMSO was added to each well. After gentle mixing, the absorbance was measured using an ELISA plate reader at 570 nm wavelength. Only DMSO was used as the blank (medium) control. For each drug, the concentration that caused only a 10% reduction in metabolic activity was used in subsequent experiments.

### 4.6. Cell transient transfection

The plasmids for EPS15 wild-type WT (Eps15-WT) and dominant negative DN Δ95/295 (Eps15-DN) constructs, both containing proteins fused to GFP ^61,62^, were kindly provided by Dra. Nathalie Sauvonnet (Institut Pasteur, Paris, France). In colocalization experiments, early and late endosomes markers, Rab5-GFP and Rab7-GFP, respectively, were used. These plasmids were a kind gift from Dr. Jose Luis Daniotti and Dr. Alejandro Vilcaes (CIQUIBIC-Universidad Nacional de Córdoba, Córdoba, Argentina). Vero cells grown to 70% confluence were transiently transfected using polyethylenimine (PEI). In brief, 2 ug of plasmid were mixed in 125 ul of ClNa 150 mM and 3 ul of PEI in DMEM. After 24 h of culture, fluorescence was observed under an epifluorescence microscope (Olympus IX81 epifluorescence inverted microscope). To evaluate the functionality of the constructs, uptake studies with TRITC-transferrin were performed on Vero cells transfected with plasmids expressing Eps15-WT or Eps15-DN, following the same procedure as described in 4.4. Transfected cell cultures were then infected with PIXV or RNV at MOI 10. Percentage of infection was calculated by scoring the number of cells positive for the viral antigen by immunofluorescence in approximately 300 transfected cells with comparable levels of GFP expression (15 microscopy fields).

In colocalization experiments of either PIXV or RNV with Rab5 and Rab7, images were acquired in a laser-scanning confocal microscope (Zeiss LSM 800 – Germany). For quantitative analysis, a single plane from z-stack cross sections images was deconvoluted using FIJI software ^63^. For each cell, a focal plane with well-defined endomembrane structures was selected for analysis and colocalization was quantified using Pearson’s correlation coefficient ^42^. Coefficient values were then used to perform the statistical analysis.

### 4.7. Inmunofluorescence assay

We performed a previously described assay ^37^. Briefly, cell monolayers were grown at 70% confluence on glass coverslips, washed three times with PBS, fixed with 4% paraformaldehyde/1% sucrose (Riedel-de Haën, Sigma-Aldrich Laborchemikalien GmbH, Seelze, Germany) for 20 min at room temperature (RT) and then washed with PBS. The cells were then permeabilized with 0.2% TritonTM X-100 (Sigma-Aldrich, St. Louis, MO) in PBS for 5 min at RT and incubated with 5% bovine serum albumin (BSA, Sigma-Aldrich, St. Louis, MO) for 1h at RT. In order to detect viral structural proteins, cells were incubated with an anti-PIXV-RNV primary polyclonal IgG antibody generated in mice in our laboratory as previously described ^37^, diluted 1/1000 in 1% BSA/PBS solution overnight at 4 °C. Then, monolayers were washed three times with PBS, and incubated with the secondary antibodies (goat anti-mouse Alexa Fluor 488- # A-11017 and Alexa Fluor 568- # A-11031, Thermo Fisher Scientific Inc., Waltham, Massachusetts, USA) for 2h at RT. To visualize the actin filaments, the fixed and permeabilized cell monolayers were labeled with Phalloidin-Tetramethyl rhodamine B (Sigma-Aldrich, St. Louis, MO) for 1h at RT. Cells were visualized using a conventional inverted epifluorescence microscope (Olympus IX81, Olympus Corporation, Shinjuku, Tokyo). Images were processed using Adobe Photoshop CS6 (Version: 13.0). Percentage of infected cells was calculated as the number of cells positive for the viral antigen from 10 microscope fields, selected at random, with comparable numbers of cells (more than 300 cells) per condition in each independent experiment. The percentage of infected cells in each treatment was calculated by normalizing data against controls (100% value).

### 4.8. Statistical analysis

All tests and graphs were performed using GraphPad Prism 5. Data are representative of at least three independent experiments and values are given as mean ± standard deviation (SD). Statistical analyses were carried out using unpaired student *t*-test and one-way ANOVA test, followed by a Tukey post-hoc analysis to enable specific group comparisons (Type I error set at 0.05), as appropriate.

## 5. Data Availability

The datasets generated during and/or analysed during the current study are available from the corresponding author on reasonable request.

## 7. Acknowledgements

We would like to thank Laura Fozzatti, Javier Aguilar, Brenda Konigheim, Guillermo Albrieu-Llinás and Pablo Lopez for generously providing some reagents; Gonzalo Quasollo and Cecilia Sampedro for technical assistance in confocal microscopy at the Centro de Micro y Nanoscopía de Córdoba, CEMINCO-CONICET-Universidad Nacional de Córdoba, Córdoba, Argentina. We particularly thank to Srta. Stella Ascheri for general technical assistance. This work was supported by grants from Secretaria de Ciencia y Tecnología - Universidad Nacional de Córdoba (SECYT-Consolidar-C N° 33620180100091CB), Ministerio de Ciencia, Tecnología e Innovación, Gobierno de Córdoba (GRFT 77/19), and Agencia Nacional de Promoción Científica y Tecnológica Argentina (FONCYT PICT/18-02324).

## 8. Author contribution

All authors were responsible for the design of the study. LMG, PIG, POQ and EG performed the experimental procedures and collected data. LMG, PIG and FMP conducted statistical data analysis. LMG, PIG and PK performed imaging analysis. PK, MC and MGP contributed ideas and comments. LMG, PIG, MC and MGP interpreted the findings and drafted the manuscript. Supervision and project administration were done by MGP and funding’s acquisition by MGP, LMG and PIG. All authors critically reviewed the contents and approved the final version for publication.

---

**Supplementary Figure S1:**
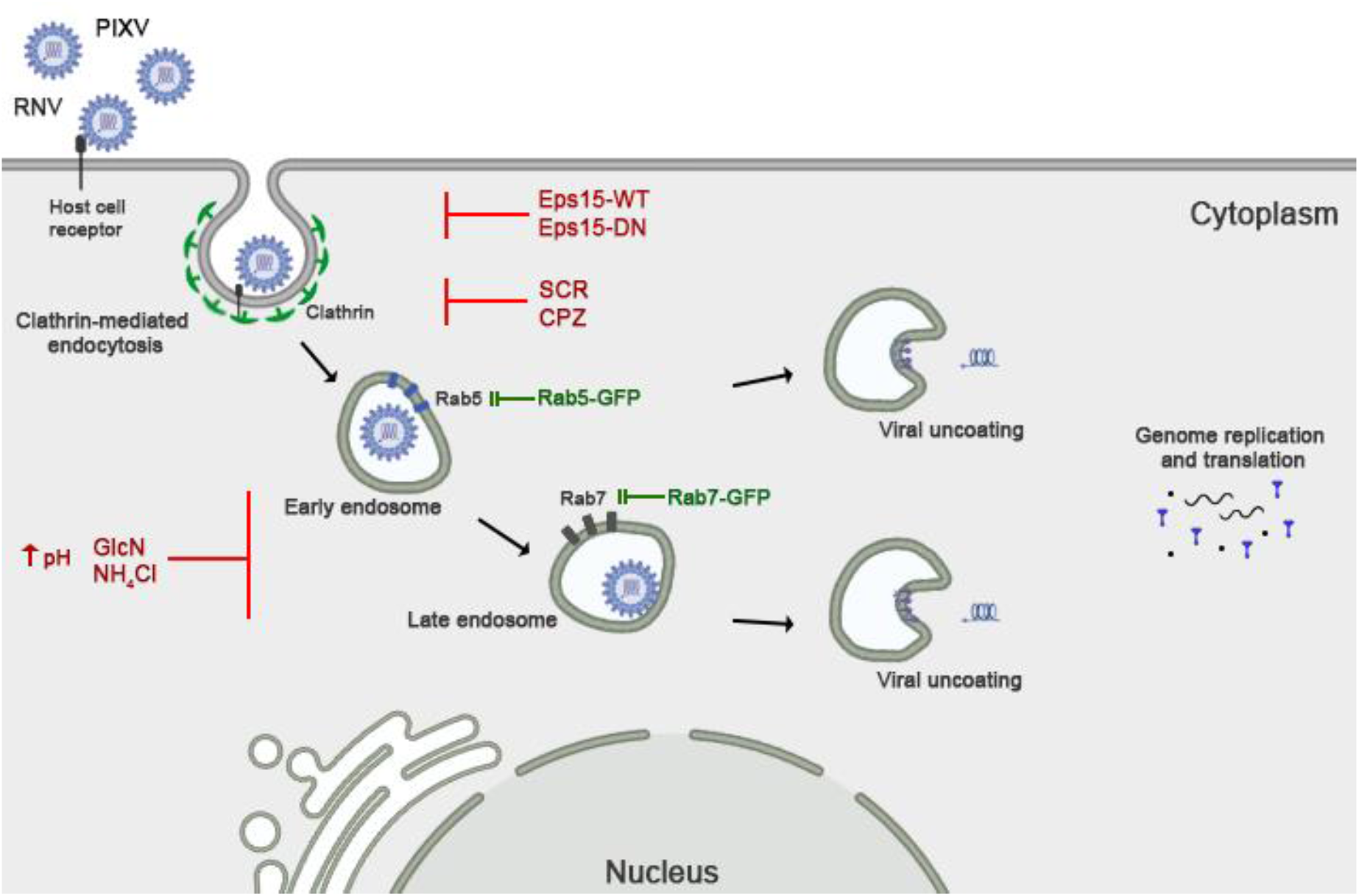
Pixuna and Rio Negro virus, members of Venezuelan Equine Encephalitis Complex, entry into host cells by clathrin-mediated endocytosis in a pH-dependent manner. **PIXV**: Pixuna virus. **RNV**: Rio Negro virus. **Eps15-WT**: Eps15 wild-type construct. **Eps15-DN**: dominant negative DN Δ95/295 mutant construct. **SCR**: Sucrose. **CPZ**: Chlorpromazine. **Rab5-GFP**: early endosomal marker. **Rab7-GFP**: late endosomal marker. **GlcN**: Glucosamine. **NH_4_Cl**: ammonium chloride.

